# Urban microbial communities shift towards carotenoid-producing species under constant anthropogenic stress

**DOI:** 10.1101/2024.11.20.624583

**Authors:** V. V. Sedova, K. A. Demin, M. P. Kulikov, E.V. Prazdnova, A.V. Gorovtsov

## Abstract

During years of investigating soil microbial communities, our research group consistently observed a predominance of carotenoid-producing microbes – characteristically pigmented in hues typical of carotenoids – in urban soil samples, as compared to those from recreational, rural, or non-urban areas. To validate this observation as non-random and to explore the ecological role of carotenoid-producing microorganisms in urban soils, we conducted a series of experiments. Additionally, we extended our study to other components of the urban ecosystem that may harbor and transfer microbial communities, including dust and snow samples. Through ecological monitoring, we assessed the proportion of culturable carotenoid-producing bacteria in those samples. It was found that in urban soils >60% of the isolates are pigmented, and in natural soils only about 30% exhibit carotenoid pigmentation. For snow and dust samples, all sites have shown pigmented bacteria predominance. Further, we analyzed both the physiological traits of individual isolates and the broader characteristics of entire culturomes (comprising all isolates from a single aliquot plated onto Petri dishes). Metagenomic analysis of the culturome washes was also performed to provide deeper insights into the microbial communities present. Metagenome of urban isolates clearly show elevated biodiversity of carotenoid biosynthesis genes and carotenoid-producing taxa, as compared to metagenome of natural habitat. In this study, *Flavobacterium* was identified as the most abundant pigmented genus of urban settings.

Based on data obtained in this work, and similar literature data on urban microbial communities, we propose that the unique combination of environmental factors in urban settings fosters the selective enrichment of carotenoid-producing microorganisms. Finally, we offer a model to explain this selective process.

## Introduction

Urban environments differ significantly from natural habitats due to the pervasive influence of anthropogenic activities. Two primary factors of urbanization that impact the environment are the accumulation of pollutants and alterations in local microclimates (Civerolo et al., 2007; Ash et al., 2008). Common urban pollutants, such as heavy metals (HMs) and polycyclic aromatic hydrocarbons (PAHs), accumulate in soils as a result of emissions from vehicular traffic and industrial activities (Liu et al., 2021; Olkova et al., 2016). Urban soils frequently exhibit disturbed structures due to mechanical compaction, which impairs gas exchange and reduces permeability. Soil pH is also often modified, typically shifting towards more alkaline conditions due to the deposition of lime, cement, and the use of saline solutions for road treatments during winter (Kurkina et al., 2012).

The lower density of vegetation in urban areas, compared to natural habitats, exacerbates soil depletion and desiccation (Ayangbenro & Babalola, 2021), further contributing to compaction and the “urban heat island” effect. This phenomenon arises from human industrial and residential activities, as well as the thermal properties of roads and buildings. As a result, urban areas are characterized by greater temperature variability, with higher minimum, average, and maximum temperatures than rural environments (McLean et al., 2005). These temperature fluctuations, coupled with seasonal freeze-thaw cycles, can alter the physical properties of soil and affect biological activity (Henry, 2007). Moreover, significant contamination with hydrophobic substances disrupts the air-thermal and water regimes of soils, reducing water availability and accelerating desiccation (Kazantseva, 2015).

Research indicates that urban conditions can affect precipitation patterns within city boundaries. Areas near major industrial sites tend to experience increased precipitation in conjunction with elevated air pollution levels (Goudie, 2018). Road traffic is a significant contributor to heavy metal and phosphorus contamination in urban snow (Viklander, 1999), while soil dust composition heavily influences rainwater quality (Feng et al., 2001). Urban dust consists of complex mixtures of pollutants, with variations depending on location and source. Anthropogenic sources of dust include vehicle emissions, fossil fuel combustion, industrial processes, and construction activities. This anthropogenic dust mixes with natural dust from soil erosion (Mihankhah et al., 2020). The movement of pollutants and the continuous circulation of dust have significant ecological implications (Deguillaume et al., 2008). For instance, polycyclic aromatic hydrocarbons (PAHs) can be transported to areas far from known pollution sources (Björseth et al., 1979). Air masses accumulate pollutants circulating over industrial areas and redistribute them over wide regions. In addition to pollutants, microorganisms can also be transported over long distances via precipitation or wind (Bauer et al., 2002). Urban pollutants, together with dust, settle on snow cover. Snow accumulates pollutants both during precipitation and after its formation, making urban snow a reliable indicator of atmospheric pollution (Engelhard et al., 2007).

These environmental conditions impose significant physiological and biochemical stress on urban microbiota. One specific stressor is desiccation; low water activity can disrupt normal protein function and compromise membrane integrity. This disruption affects the electron transport chain, leading to the accumulation of free radicals, protein denaturation, DNA mutations, and lipid peroxidation (Ayangbenro & Babalola, 2021). The effects of desiccation are exacerbated by UV radiation and heavy metal pollution. Oxidative metals, such as copper, chromium, iron, and vanadium, participate in reactions analogous to Fenton reactions, generating reactive oxygen species. Non-oxidative metals, including arsenic, cadmium, mercury, nickel, lead, and antimony, can deactivate antioxidant enzymes, disrupting the balance between oxidants and antioxidants (Hobman & Crossman, 2015). The overall degradation of urban soils leads to reduced plant cover, and when combined with the removal of leaf litter, this diminishes organic matter content, potentially slowing microbial growth rates.

The study of microbial community transformations in response to urbanization is a relatively recent field of inquiry. Nevertheless, numerous studies have demonstrated that urbanization significantly affects microbial community composition. An increase in the diversity of Pseudomonadota, Actinomycetota, and Bacteroidota, alongside a decrease in Nitrospirota and Gemmatimonadota, has been reported in the works of Li et al. (2023) and Whitehead et al. (2022). Hu et al. (2018) showed that high compaction in urban soils alters the composition of soil communities, leading to a decline in overall diversity. In urban soils, genera such as *Arthrobacter, Sphingomonas*, and *Chryseolinea* predominate. Interestingly, all three genera, despite belonging to different phyla, are carotenoid producers, and their colonies are typically yellow-pigmented. A reduction in overall microbial activity in response to soil compaction has also been noted (Li et al., 2023).

Several studies have examined the bacterial composition of urban precipitation. In urban environments, characterized by high dust volumes carrying microorganisms over considerable distances, these microbes are exposed to various stressors, including UV radiation and desiccation under oligotrophic conditions (Meola et al., 2015). Pollutants within dust can further adversely affect microbial cells (Nikolaeva et al., 2022). Dust aerosols have been found to be dominated by representatives of the genera *Micrococcus, Myxococcus*, and *Cytophaga* (Lysak et al., 2023), which also produce yellow carotenoids. It has also been shown that, in urban environments, the diversity of airborne bacteria is lower than in natural environments, while their overall concentrations are higher (Després et al., 2012).

It has been shown that both old and fresh snow harbour a fairly high diversity of microorganisms. Dominant groups include Bacteroidota (39%), molds (27%), coccoid microorganisms (24.2%), and bacilli (9.4%) (Lykov, Morozova, & Krymskaya, 2021). Additionally, there is a notable prevalence of pigment-producing bacteria, including representatives of *Pseudomonas, Micrococcus*, and *Rhodococcus* (Naprasnikova & Makarova, 2006).

In summary, various factors characteristic of urban environments—such as pollutants, altered microclimates, and physical stressors—substantially influence microbial life. A common outcome is the emergence of oxidative stress and reduced metabolic activity, which favour more resilient species. Over time, this leads to structural changes in microbial communities. One notable shift is the increased abundance of species that produce carotenoids, which are known for their antioxidant, anti-UV, and anti-desiccation properties. As we and other researchers have observed, this shift appears to be a recurring phenomenon in urban microbial communities. Thus, the goal of our research was to investigate the ecological mechanisms underlying this trend.

## Materials and methods

### Sampling sites description

The overall analysis was conducted using samples collected from the city of Rostov-on-Don (southern Russia) and its surrounding areas. To assess the level of urbanization at the sampling sites, a combination of factors was considered. Firstly, for a site to be classified as having low urbanization, it needed to have abundant vegetation cover, primarily grasses or shrubs (areas under dense tree cover were avoided to exclude the influence of trees as a significant factor affecting the underlying soil microbial community). Secondly, there should be no presence of technogenic materials, such as paper, metal, glass debris, construction waste, or plastic litter. Thirdly, the site had to be located far from urban infrastructure, roadways, or other urban developments, and thus characterized by a low proportion of sealed surfaces. Sites that did not meet these criteria were classified as urbanized. In the case of urbanized soils, these sites were typically characterized by high levels of compaction and surface sealing, or proximity to roadways. Additionally, data from a three-year monitoring programme on dust deposition across various districts of Rostov-on-Don and adjacent areas were considered, including the mass and composition of dust and solid particles deposited per km^2^ (Privalenko & Bezuglova, 2003). Finally, a water repellency test (WDPT test) was performed on each soil sample as an additional criterion for classification (Korenkova & Matush, 2015).

### Sampling of soil, snow, and dust material

Soil samples were collected according to ISO 10381-1 (2002). The top 0–5 cm layer was sampled using the envelope method and placed in sterile plastic bags. Mixed samples from each site, weighing approximately 1.5 kg, were transported to the laboratory in a cooled state and stored at 4°C for no longer than 7 days before further analysis. During the winter months (January-February), snow samples of approximately 10 cm^3^ were collected from the same sites using sterile plastic containers. Dust samples were collected from large, exposed surfaces, such as stone, asphalt, or anthropogenic objects (free of moss or lichen colonization), including windowsills, metal structures, and similar surfaces. Dust sampling was performed as follows: the sampling surface was divided into several segments, and each segment was wiped with a pre-weighed sterile gauze cloth. These cloth fragments were then placed in sterile containers.

Soil sampling was conducted across three seasons: autumn, winter, and spring. Snow samples were collected in winter, and dust samples were collected in autumn.

### Plating experiments and colonies enumeration

For soil samples, 10 g of soil material, previously homogenized in a porcelain mortar with a rubber pestle, was added to a flask containing 100 ml of sterile tap water, after which serial dilutions were performed up to 10^−4^. For snow samples, one milliliter of melted snow water was diluted until no solid suspended particles were observed in the dilution. For dust samples, the gauze cloth fragments with dust material were placed in a flask, and shacked on a rotary shaker for 30 minutes to allow dust particles and cells transfer into water suspension.

To determine the optimal working dilution for the inoculation and enumeration of cultivable microorganisms from dust, snow, and soil samples, we conducted a preliminary experiment. Serial dilutions of each sample were plated onto a nutrient medium, and the dilution yielding 30–300 colonies after 7 days of incubation was identified. For the main experiments, 50 μl aliquots of the working dilution were inoculated onto the surface of the medium and evenly spread with a sterile spatula until the inoculum was fully absorbed. The working dilutions used were 1:10,000 for soil, typically 1:1,000 for dust, and typically 1:100 for snow. Inoculated plates were incubated for 7 days at the appropriate conditions.

The nutrient medium used in the experiments was soy agar, which contained the following components per liter: soy flour, 25 g; NaCl, 1.5 g; glucose, 1.5 g; CaCOLJ, 1.5 g; and agar, 25 g.

Each sampling site was represented by two biological replicates, with each biological replicate consisting of three analytical replicates (i.e., one sampling site = two independent sample dilutions = six plates). After the incubation period, photographs of all plates were taken. Colony counts were performed using Adobe Photoshop 2022, employing the “Counter” tool. For each plate, the total number of colonies was recorded, along with the number of colonies exhibiting yellow, red, orange, or pink pigmentation.

Colonies morphologically resembling Streptomyces spp.—characterized by pigmentation ranging from orange to brown to black, or by the production of similar pigments diffused into the medium—were excluded from the count of colored colonies. This exclusion was implemented to avoid misclassification of melanin-producing colonies as carotenoid-producing colonies.

### Biomass collection and carotenoid extraction

After incubation and colony counting, biomass was collected from the plate surfaces. For this, 2 ml of acetone was applied to the plate surface, and the biomass was washed off using a spatula. The biomass was then transferred to a microcentrifuge tube and centrifuged for 10 minutes at 7000g, with the subsequent removal of supernatant. The microcentrifuge tubes containing the biomass were frozen for 24 hours. For extraction, after the biomass thawed, the following procedure was performed three times: 1 ml of acetone was added to the tube containing the biomass, and the cell pellet was disrupted in acetone using a Sonics VCX130 ultrasonic homogenizer; the biomass suspension in acetone was then centrifuged for 5 minutes at 5000g; the colored acetone supernatant was transferred to a clean microcentrifuge tube.

Thus, for each wash from the plate surface, three colored fractions were combined for a total volume of 3 ml (hereafter referred to as the working extract). A total of 200 μl of the working extract from each sample was added to the wells of a polystyrene 96-well microplate. Each sample was represented by three working extracts (obtained from biomass from three plates). The microplates with extracts were analyzed using the Fluostar Omega plate reader. The absorption spectrum of each working extract was measured in the wavelength range from 250 to 950 nm. The resulting OD (optical density) values for each wavelength were used for further analysis.

### Microbial species’ isolation and identification

Microbial species were isolated using tryptic soy agar (TSA) medium. Taxonomic identification was carried out in Scryabin Institute of Biochemistry and Physiology of Microorganisms (Pushchino, Russia) by sequencing V3-V4 16s rRNA gene. To obtain absorption spectra of isolates’ biomass, the same method as described before was used.

### Inhibition of carotenoid synthesis and catalase test

To assess whether carotenoids are responsible for isolates’ stress tolerance, we chemically inhibited carotenoid biosynthesis. This chemical knock-out was achieved by adding diphenylamine (DPA) to the solid nutrient medium (Salton, Schmitt, 1967). The DPA concentration was individually adjusted for each isolate. The final concentration range at which carotenoid synthesis ceased, but the culture growth pattern did not differ from the control without DPA, was 100–200 μM.

To analyze the presence of catalase, a hydrogen peroxide test was used (Ahmad et al., 2014). Bacterial cultures were applied to a microscope slide, and one drop of 3% H2O2 solution was added. The formation of bubbles and foam within 5–10 seconds after adding the peroxide was used as a positive criterion for the presence of the enzyme. The analyses were repeated three times.

### Determination of H2O2 Resistance

The biomass of a two-day culture of the control strain (incubated on soy agar) and the strain variant with inhibited carotenoid synthesis ability (incubated on soy agar + DPA) was suspended in a phosphate buffer and diluted to a cell density of 1 McFarland. This 1 McF suspension was divided into 4 parts. Hydrogen peroxide was added to each part at concentrations of 0, 5, 20, and 50 mM respectively, and incubated for 1 hour. Subsequently, serial dilutions (up to 10^3^) from each variant were plated. The plates were incubated for 5 days. At the end of the incubation, colonies were enumerated using the dilution which yielded CFU in the range of 30-300. Strains and their variants which were unable to produce pigments were compared.

### Metagenomic analyses

#### Experiment design

To characterize carotenoid biosynthesis pathways in microorganisms from urban and non-urban sites, extraction and sequencing of the total DNA from soil culturomes was performed. Two contrasting sample sites were used to obtain representative culturomes: the site in the city center (Voroshilovskiy av., 47.2320, 39.7143) and the site located on the territory of Rostov-on-Don botanical garden (47.2373, 39.6584). Soil samples were plated as described above in two biological and three analytical replicates (6 plates per sample). After 10 days of incubation, biomass from 6 plates was washed with sterile distilled water, centrifuged for 10 minutes at 10000g and collected into one sample. Two samples, further referred as BS (botanical garden) and VK (city center), were subjected to DNA extraction. DNA was isolated using DNA-Extran kit (Syntol, Russia) according to the instructions. Sequencing was performed on a MinION device using the Rapid Sequencing Kit V14 reagent kit according to the instructions.

#### Bioinformatics

Quality control of the reads was performed using NanoPlot v1.42.0 (De Coster and Rademakers et al., 2023), filtering and discarding of low-quality reads was performed using Filtlong v0.2.1. Reads were assembled into contigs using MEGAHIT (Li, Dinghua et al., 2015). Each contig was taxonomically classified based on its ORF predicted by prodigal. Briefly, the CATpack program (Hauptfeld et al., 2024) was used to align predicted ORFs with the GTDB reference database of prokaryotic proteins. Only contigs classified on the basis of at least two ORFs were retained for the analysis. To annotate and functionally characterize obtained contigs, EggNOG mapper software (Cantalapiedra et al., 2021) with mmseqs database was used. We specifically searched the genes coding for components of carotenoid biosynthesis pathways and xenobiotics degradations/tolerance pathways (complete list is presented in the supplementary table 1).

## Results

### Ecological monitoring of culturable carotenoid-producing microbial communities

During all the experiments, we observed an increased share of carotenoid-producing colonies from urban sites. The isolates of urban soils were characterized by a wide variety of carotenoid-associated colors: yellow, orange, red, coral and pink. Colonies of pink and red color were the least frequently encountered, typically associated with cultures derived from snow and dust samples. Among natural soils, non-pigmented colonies, as well as actinomycetes-like and bacilli-like colonies were more commonly isolated, while pigmented forms were represented in smaller numbers and exhibited less diversity, primarily displaying various shades of yellow. Figure 1 (a, b, c) displays the percentages of pigmented colonies when plating samples from urbanized and non-urbanized habitats (soil, dust, snow).

**Figure 1.**
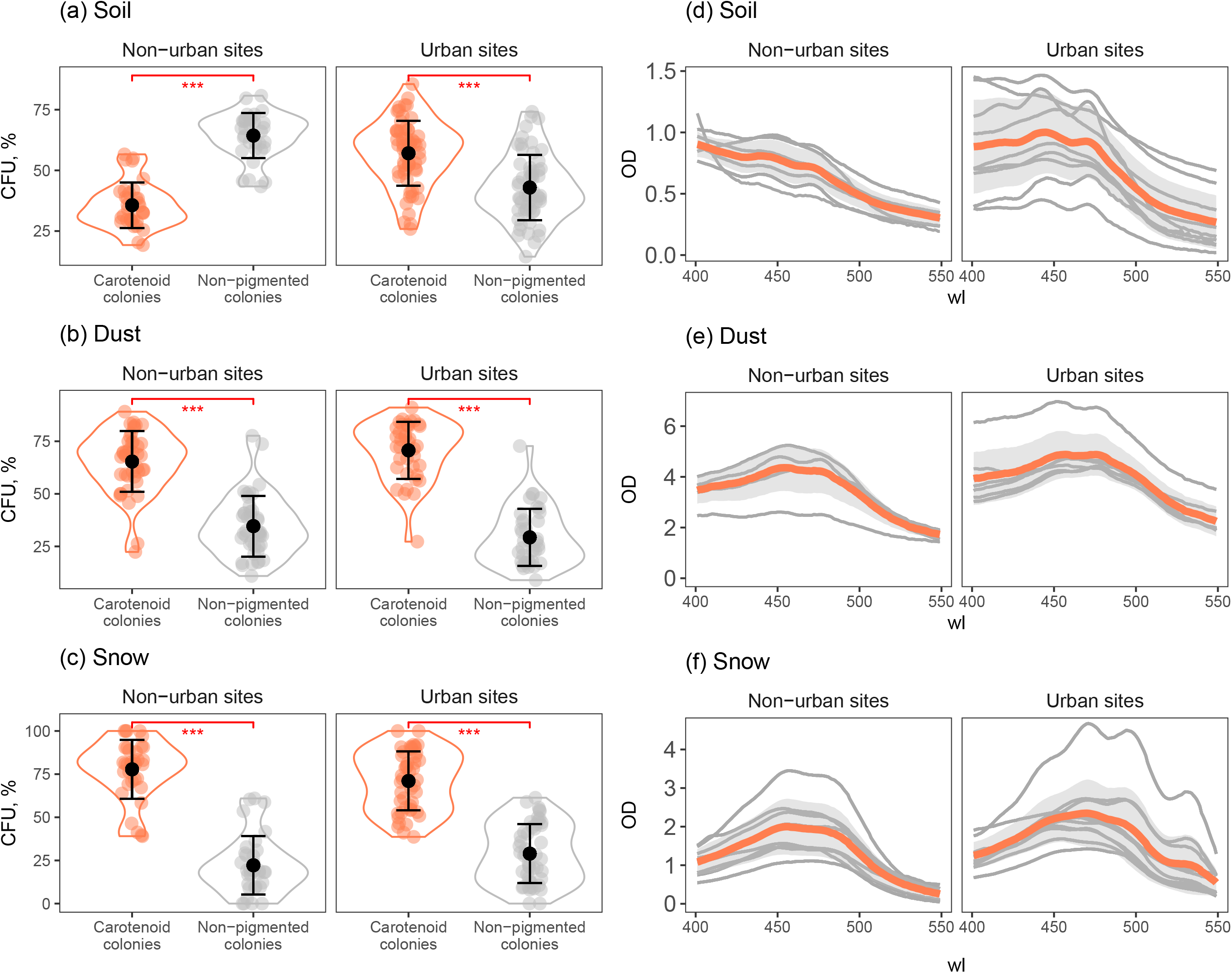
Facets **a, b, c**: the abundance of carotenoid-pigmented colonies and non-pigmented colonies isolated from urban and non-urban soil (**a**), dust (**b**) and snow (**c**) material. Facets **d, e, f**: absorption pics of microbial biomass isolated from urban and non-urban soil (**d**), dust (**e**) and snow (**f**) material. Gray lines represent different sampling sites, coral lines represent an average curve for the area.

Urban and non-urban soils differ significantly in the number of pigmented colonies (*p* <0.0001, Mann-Whitney U-test). For soils collected in the city, the average proportion of carotenoid-producing CFUs is 61% ± 13, whereas for soils with low urbanization levels, it is 34% ± 9. Among dust and snow samples, the difference between urbanized and non-urbanized territories was less pronounced or not observed: for urban dust deposits, pigmented CFU reached 70±13%, while for sites outside the city – 65±15% (*p* = 0.04). For urban snow deposits – 71±17%, outside the city – 79±17% (*p* = 0.08). Abundance of non-pigmented colonies was higher than of pigmented ones only for non-urban soil samples, while for soil urban samples, as well as for all snow and dust and samples, pigmented colonies abundances was always the highest (for each comparison, *p* < 0.0001).

The absorption spectra of biomass grown from soil, snow, and dust samples were also analyzed (Figure 1d, e, f). The results of the light absorption analysis were generally in accordance with the data obtained from plates.

For snow and dust samples, carotenoid-specific peaks are clearly distinguishable for both urban and non-urban areas. However, the absorption spectra of extracts from snow biomass were more shifted towards the red region of the spectrum for all samples compared to similar data for soils and dust.

The absence of clear difference between urban snow/dust spectra and non-urban snow/dust spectra is highly likely explained by the transfer of dust particles and pigmented cells far beyond the location of pollution sources by air masses. At the same time, even despite the redistribution of solid particles and cells raised into the air, the effect remains more pronounced for urban areas. For example, the biomass isolated from snow was characterized by a greater diversity in wavelengths. Absorption peaks at 530-545 nm are visible for samples from urban sites. It is known that some carotenoids can absorb near 530 nm, such as saproxanthin (Planctomycetota, *Rhodopirellula* sp.), diapolycopene (Bacillota, *Heliobacterium* sp.), and bacterioruberin (Archaea, *Halobacterium* spp.). In soils of natural zones, such peaks are absent.

### Pigmented isolates testing

During our experiments, we have been isolating microorganisms representing different colors of carotenoid pigments. We reached several hundred isolates and tried to maintain as many of them as possible in culture. However, only a small fraction survived over several repetitive transfers. We were able to assess colors ratio, as well as identify the taxonomy of several morphologies which were encountered the most frequently and thus were representative of the group. Finally, we tested the oxygen stress tolerance in the representative isolates (Figure 2a,b,c,d).

**Figure 2.**
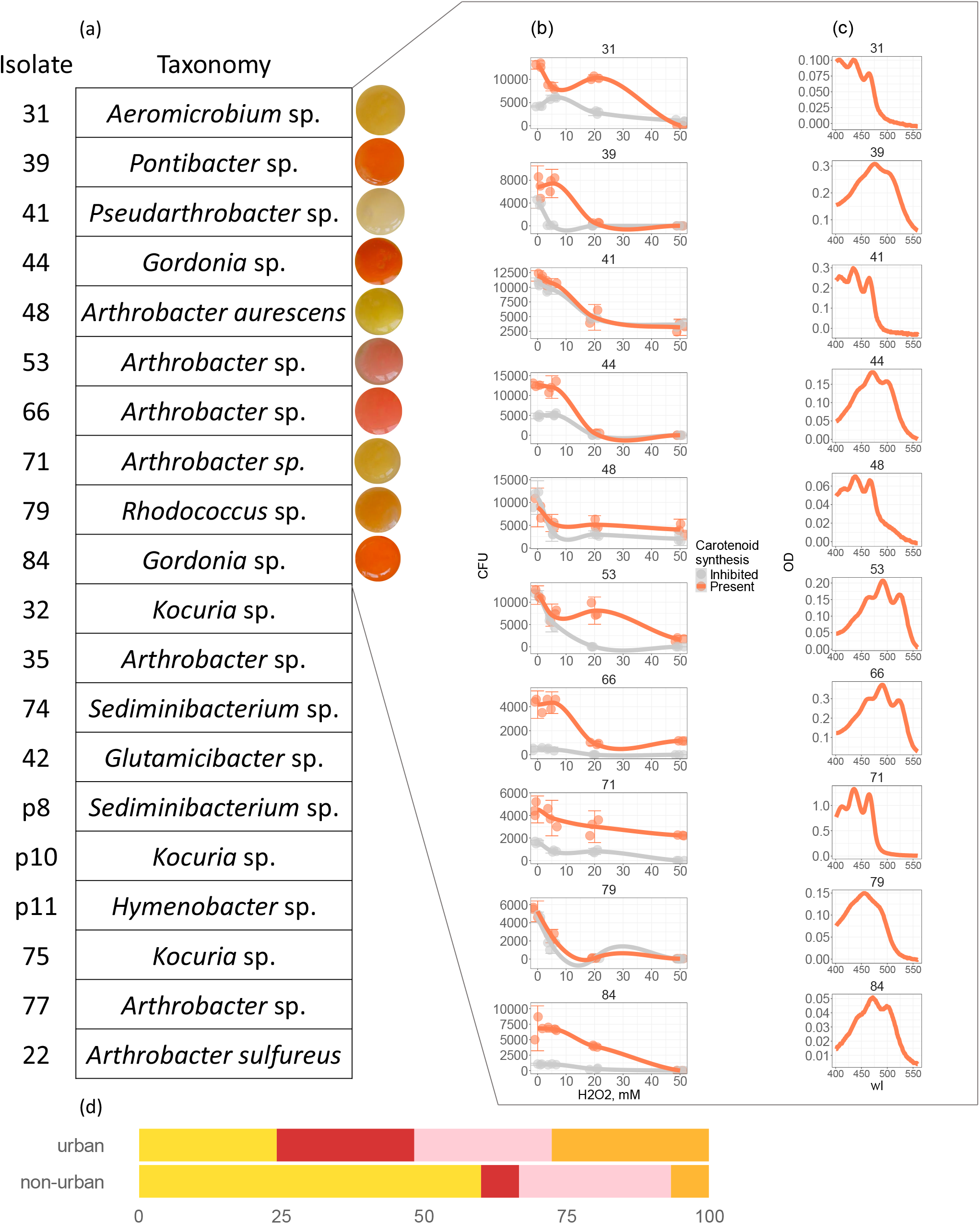
Facet **a**: taxonomy of different isolates obtained in this study. All the species isolated from urban soil samples. Facet **b, c**: H2O2 tolerance test performed on ten isolates and their variants chemically knocked-out for the ability to produce carotenoids (**b**); absorption spectra obtained from isolates biomass (**c**). Facet **d**: overall abundance in % of four major hues exhibited by microbial colonies from natural and urban areas.

Isolates from urban soil samples consistently exhibited much more color diversity than that of non-urban sites, with pink and red colonies relative abundance near ∼25% each over the several years of monitoring. Non-urban sites usually isolate yellow-pigmented bacteria (Figure 2d).

The taxonomic position was obtained for 20 isolates. Classified pigment-producing isolates were affiliated with microbial genera consistently observed in the context of anthropogenic pollution or urban microbiota studies. Most isolates were represented by carotenoid-producing Actinomycetota genera *Arthrobacter, Pseudoarthrobacter, Gordonia, Kocuria* and *Glutamicibacter*. The rest majority was of Bacteroidota phylum, particularly representing Chitinophagales (*Sediminibacterium*) and Cythophagales (*Pontibacter, Hymenobacter*) orders. Most of the tested strains were able to tolerate 20 mM of hydrogen peroxide, while only isolates №48 (*Arthrobacter aurescens*), 41 (*Paenarthrobacter* sp.), №66 and №71 (*Arthrobacter* spp.) survived over 1 hour of 50 mM treatment. Strikingly, isolates №48 and №41 tolerated 50 mM of H2O2 even with inhibited carotenoid biosynthesis. *Paenarthrobacter* isolate did not differ between its wild and chemically knocked-out form in the ability to tolerate reactive oxygen species, which may be attributed either to the firm cell wall and overexpressed enzymatic mechanisms (e.g., catalase, SOD). Notably, two of all *Arthrobacter* isolates produced red carotenoids independently of oxygen stress level, which is a rare feature for this mostly yellowish group of actinobacteria. Their absorption peaks were – 465, 490, 522 nm for isolate №66 and 460, 491, 524 nm for isolate №53.

### Metagenomic analysis of urban and non-urban culturomes

In our survey, we hypothesized that metagenomic data from urban soils would exhibit elevated copy numbers and greater biodiversity of carotenoid synthesis genes. The results partially supported this expectation. Contigs assembled from urban soil metagenomes were affiliated with a diverse range of taxa, including *Flavobacterium, Pedobacter, Arthrobacter, Paenarthrobacter, Sphingobium, Novosphingobium, Dyadobacter, Epilithonimonas*, and *Chitinophaga*. These findings are consistent with the hypothesis that urban soils harbor a high diversity of carotenoid-producing bacteria.

In contrast, contigs assembled from non-urbanized sites were predominantly associated with aerobic spore-forming bacteria belonging to the Bacillota phylum, including genera such as *Bacillus, Paenibacillus, Brevibacillus*, and *Priestia* (Figure 3c).

**Figure 3.**
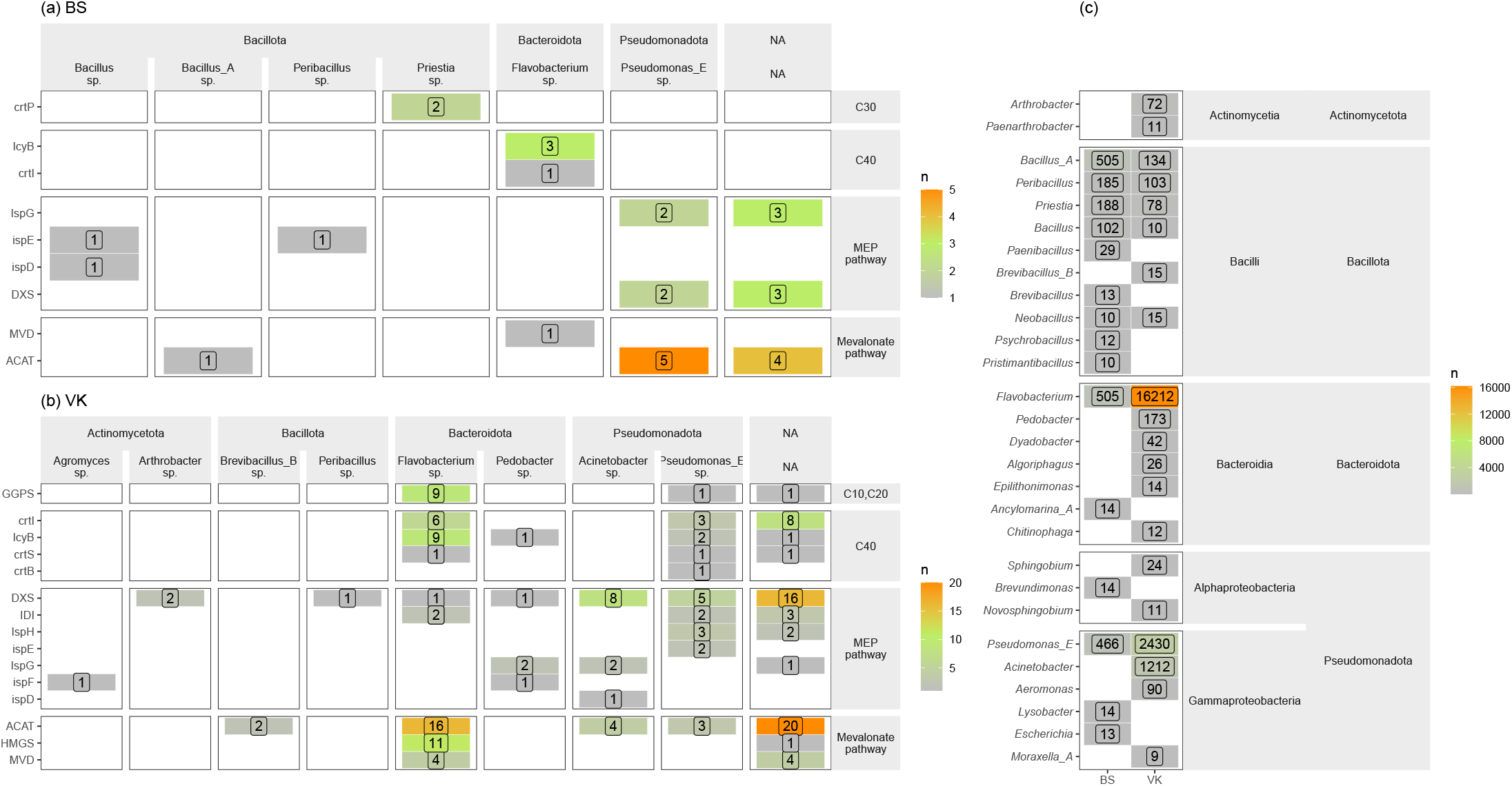
Facets **a, b**: taxonomy, as well as abundance of carotenoid biosynthesis genes and genes required for carotenoid precursor synthesis identified in the metagenomic contigs of natural (**a**) and urban (**b**) areas. Facet **c:** Bulk contigs abundance and their taxonomic affiliation in two metagenomic samples.

Moreover, metagenomes from the urban soil sample were characterized by an increased abundance of carotenoid biosynthesis genes (Figure 3a,b). While multiple pigment-producing genera were identified at the bulk contig level, the majority of carotenoid biosynthesis genes were primarily affiliated with the genera *Flavobacterium* and *Pseudomonas*. A smaller fraction of terpenoid backbone biosynthesis genes was associated with contigs from *Pedobacter* and *Arthrobacter*. Additionally, some contigs in the urban metagenome could not be taxonomically assigned but encoded genes involved in carotenoid, mevalonate, and terpenoid biosynthesis pathways.

Non-urban metagenome contained the *crtP* gene, encoding diapolycopene oxygenase, within *Priestia* contigs. The *lcyB* and *crtI* genes, associated with the biosynthesis of C40 carotenoids, were identified in *Flavobacterium* contigs.

Overall, contigs in the urban metagenome encoded the the synthesis of a wide range of carotenoids, including phytofluene, neurosporene, lycopene, γ-carotene, ζ-carotene, β-carotene, α-carotene, zeaxanthin, cryptoxanthin, adonixanthin, zeinoxanthin, lutein, astaxanthin, canthaxanthin, echinenone, β-zeacarotene, and dihydro-β-carotene. In contrast, contigs from the non-urban site metagenome harbored *crtP*, indicative of an incomplete staphyloxanthin biosynthesis pathway, along with genes required for the biosynthesis of phytofluene, neurosporene, lycopene, γ-carotene, ζ-carotene, β-carotene, β-zeacarotene, and dihydro-β-carotene.

## Discussion

Based on the results, we infer that carotenoid-synthesizing bacteria play a significant role in urban habitats, as demonstrated by analyses of urban soil, dust, and snow precipitation. An increase in the proportion of pigmented bacterial forms was specifically observed in urban soil samples, whereas this trend was not evident in the analyzed dust and snow samples. It is hypothesized that urban conditions may favor the selection of more resilient microbial groups, particularly those capable of synthesizing carotenoids.

Soil particles, along with their associated microorganisms, are transported by wind from exposed urban soils, contributing to the dust that circulates in the atmosphere. This process facilitates the long-distance transport of microorganisms, which are subsequently exposed to a variety of environmental stressors, including ultraviolet (UV) radiation, oligotrophic conditions, pollutants present in the dust, desiccation, and disruptions in microbe-microbe interactions. These factors likely contribute to the selection of more resilient microbial groups, which then settle and accumulate in the surface layers of urban soils.

In contrast, in areas with low levels of urbanization, such adaptations may not offer a competitive advantage, allowing these microbial groups to be supplanted by more stable, established communities. However, within urban soils—where many of the aforementioned stressors are persistent—carotenoid-producing microorganisms may acquire a selective advantage.

Metagenomic data and our experiments with isolated cultures are in agreement with this model. We observe that the ability to tolerate oxygen stress is mostly attributed to the presence of carotenoids, while the specific genes involved in carotenoid metabolism are more diverse and represented in urban samples. The study of (Lysak et al., 2023) yielded similar results. Despite that, it seems that the specific microbial community composition would differ from one urban environment to another. For example, a 2022 study investigated the diversity of airborne microbial communities in contaminated areas of Rome, examining bacterial communities in road dust and on leaf surfaces. In road dust, Pseudomonadota (24%) and Bacteroidota (12%) were the most represented phyla, with many of these organisms characterized by pigmented colonies. These included *Stenotrophobacter roseus, Arenimicrobium luteum, Skermania piniformis, Kineococcus siccus, Yaniella halotolerans*, and *Rhodocytophaga rosea* (Pollegioni et al., 2022).

Snow provides even harsher conditions for microbial species selection. Microbial communities in the cryosphere have been primarily studied in high-altitude and remote areas. Snow microbial communities are known to be influenced by microorganisms introduced from other environments via dust and wind. For example, it has been suggested that dust settling on the snow of the San Juan Mountains may originate from distant regions such as the Mojave Desert or Colorado Plateau, affecting microbial adaptability through selective pressures in dusty conditions (Courville et al., 2020). In urban environments, where snow cover is seasonal, microbial circulation occurs: bacteria from soil surfaces enter the atmosphere and settle on the snow. Urban snow and ice cover, affected by repeated freeze-thaw cycles, further influence microbial communities. Urban microclimates accelerate snow melting due to the heat island effect, coupled with the use of de-icing agents. Nutrient availability is limited under these conditions, while UV exposure may be prolonged due to snow’s high refractive index (Sajjad et al., 2020). After snowmelt, microbial communities are introduced back into the soil (Naprasnikova & Makarova, 2006). Apparently, microbial mass entering the soil after such repetitive cycles of exposure to diverse stresses, are enriched with pigmented species and strains, especially red-colored, as red pigmentation is considered more protective against UV, ROS or desiccation effects (Dieser et al., 2010). It is also known that lowering the incubation temperature leads to a more reddish coloration of microbial species capable of carotenoid production (Sutthiwong et al., 2014). Here, we also see such an effect: microbial biomass absorption spectra of the snow are shifted to the red part of the visible wavelength range, while isolates from urban soil samples are pigmented red more often than those from natural habitats. Moreover, we managed to isolate red-pigmented species of *Arthrobacter* genus, while species of this genus are generally yellow (Sutthiwong et al., 2014). Recently, a novel red carotenoid compound with anticancer activity was discovered from *Arthrobacter* (Alfra et al., 2017). In the study of Reddy et al. (2003), carotenoid extracts from psychrophilic *Arthrobacter roseus* bacterium demonstrated peaks at 467, 494 and 524 nm, which is near the peaks we observed in this study for two of our *Arthrobacter* isolates (465, 490, 522 nm for isolate №66 and 460, 491, 524 nm for isolate №53). Despite the fact that *Arthrobacter* species may produce porphyrins and thus exhibit red colors, we clearly show, using diphenylamine inhibition method, that their ROS tolerance depends on carotenoid production. The testes with other isolates resulted mostly in the same conclusion.

Our metagenomic analysis, however, have shown different microbial community composition, mainly because of the restricted sample size. Sequenced culturomes have shown Pseudomonadota and Bacteroidota as the dominant pigmented species in the cultured component of urban microbial community. It is highly likely that the exact composition of carotenoid-producing bacteria would vary drastically between cities of different sizes and levels of industrialization and integration with natural components, such as squares, gardens and other recreational “green” zones. The selection, however, would be the same, with microbes undergoing selective stress if introduced to the atmosphere and precipitation from soil.

## Conclusions and future perspective

An increase in the proportion of pigmented bacteria was observed in urban soil samples, whereas this pattern was not seen in dust and snow samples. In soils collected from urban areas, the average proportion of carotenoid-forming colony-forming units (CFUs) was 61%±13, compared to 34%±9 in soils from less urbanized areas. For dust and snow samples, the difference between urbanized and non-urbanized areas was less pronounced or not observed: for urban dust deposits, it was 70%±13, compared to 65%±15 outside the city. For urban snow deposits, it was 71%±17, compared to 79%±17 outside the city. The data obtained from biomass extracts confirmed the trend observed in CFU counts.

Twenty pigmented isolates were identified, most of which being affiliated with Actinomycetota and Bacteroidota phyla, and *Arthrobacter* as the most frequently encountered genus. Ten isolates were tested for ROS stress tolerance, and their biomass was extracted to analyze the absorption peaks. Most of them survived 20 mM hydrogen peroxide treatment, but only four survived 50 mM.

Metagenomic analysis of culturomes isolated from two contrasting soil samples confirmed the observations. City center microorganisms represented increased abundance and diversity of carotenoid-producing species and carotenoid metabolism genes.

Consequently, the findings presented here provide a foundation for further investigation into carotenoid-producing bacteria within urbanized environments. This research could enhance our understanding of urban ecosystems and support the development of innovative methods for monitoring technogenically transformed soils, as well as identifying new sources for the industrial production of carotenoid pigments.

